# From shallow to deep: exploiting feature-based classifiers for domain adaptation in semantic segmentation

**DOI:** 10.1101/2021.11.09.467925

**Authors:** Alex Matskevych, Adrian Wolny, Constantin Pape, Anna Kreshuk

## Abstract

The remarkable performance of Convolutional Neural Networks on image segmentation tasks comes at the cost of a large amount of pixelwise annotated images that have to be segmented for training. In contrast, feature-based learning methods, such as the Random Forest, require little training data, but never reach the segmentation accuracy of CNNs. This work bridges the two approaches in a transfer learning setting. We show that a CNN can be trained to correct the errors of the Random Forest in the source domain and then be applied to correct such errors in the target domain without retraining, as the domain shift between the Random Forest predictions is much smaller than between the raw data. By leveraging a few brushstrokes as annotations in the target domain, the method can deliver segmentations that are sufficiently accurate to act as pseudo-labels for target-domain CNN training. We demonstrate the performance of the method on several datasets with the challenging tasks of mitochondria, membrane and nuclear segmentation. It yields excellent performance compared to microscopy domain adaptation baselines, especially when a significant domain shift is involved.

## 1 INTRODUCTION

Semantic segmentation – partitioning the image into areas of biological (semantic) meaning – is a ubiquitous problem in microscopy image analysis. Compared to natural images, microscopy segmentation problems are particularly well suited for feature-based (“shallow”) machine learning, as the difference between semantic classes can often be captured in local edge, texture or intensity descriptors (Berg et al. (2019); Arganda-Carreras et al. (2017); Belevich et al. (2016)). While convolutional neural networks (CNNs) have long overtaken feature-based approaches in segmentation accuracy and inference speed, interactive feature-based solutions continue to attract users due to the low requirements to training data volumes, nearly real-time training speeds and general simplicity of the setup, which does not require computational expertise.

CNNs are made up of millions of learnable parameters which have to be configured based on user-provided training examples. With insufficient training data, CNNs are very prone to overfitting, “memorizing” the training data instead of deriving generalizable rules. Strategies to suppress overfitting include data augmentation (Ronneberger et al. (2015)), incorporation of prior information (El Jurdi et al. (2021)), dropout and sub-network re-initialization (Taha et al. (2021); Han et al. (2016)) and, in case a similar task has already been solved on sufficiently similar data, domain adaptation and transfer learning. In the latter case, the network exploits a large amount of labels in the so called “source” domain to learn good parameter values for the task at hand, which are further adapted for the unlabeled or sparsely labeled “target” domain through unsupervised or weakly supervised learning. For microscopy images, the adaptation is commonly achieved by bringing the distributions of the source and target domain data closer to each other, either by forcing the network to learn domain-invariant features (Roels et al. (2019); Liu et al. (2020); Long et al. (2015)) or by using generative networks and cycle consistency constraints (Januszewski and Jain (2019); Zhang et al. (2018); Chen et al. (2019)). Alternatively, the domain shift can be explicitly learned in a part of the network (Rozantsev et al. (2018)). In addition to labels in the source domain, pseudo-labels in the target domain are often used for training (Choi et al. (2019); Xing et al. (2019)). Pseudo-labels can be computed from the predictions of the source domain network (Choi et al. (2019)) or predictions for pixels similar to source domain labels (Bermúdez-Chacón et al. (2019)).

In contrast, Random Forest (RF), one of the most popular “shallow” learning classifiers (Fernández-Delgado et al. (2014)), does not overfit on small amounts of training data and trains so fast that in practice no domain adaptation strategies are applied – the classifier is instead fully retrained with sparse labels in the target domain. However, unlike a CNN, it cannot fully profit from large amounts of training data. The aim of our contribution is to combine the best of both worlds, exploiting fast training of the Random Forest for domain adaptation and excellent performance of CNNs for accurate segmentation with large amounts of training data. We use the densely labeled source domain to train many Random Forests for segmentation and then train a CNN for Random Forest prediction enhancement (see Figure 1). On the target domain, we train a new Random Forest from a few brushstroke labels and simply apply the pre-trained Prediction Enhancer (PE) network to improve the probability maps. The enhanced predictions are substantially more accurate than the Random Forest or a segmentation CNN trained only on the source domain. Furthermore, a new CNN can be trained using enhanced predictions as pseudo-labels, achieving an even better accuracy with no additional annotation cost. Since the Prediction Enhancer is only trained on RF probability maps, it remains agnostic to the appearance of the raw data and can therefore be applied to mitigate even very large domain gaps between source and target datasets, as long as the segmentation task itself remains similar. To illustrate the power of our approach, we demonstrate domain adaptation between different datasets of the same modality, and also from confocal to light sheet microscopy, from electron to confocal microscopy and from fluorescent light microscopy to histology. From the user perspective, domain adaptation is realized in a straightforward, user-friendly setting of training a regular U-Net, without adversarial elements or task re-weighting. Furthermore, a well-trained Prediction Enhancer network can be used without retraining, only requiring training of the Random Forest from the user. Our Prediction Enhancer networks for mitochondria, nuclei or membrane segmentation tasks are available at the BioImage Model Zoo (https://bioimage.io) and can easily be applied to improve predictions of the Pixel Classification workflow in ilastik or of the Weka Trainable Segmentation plugin in Fiji.

**Figure 1.**
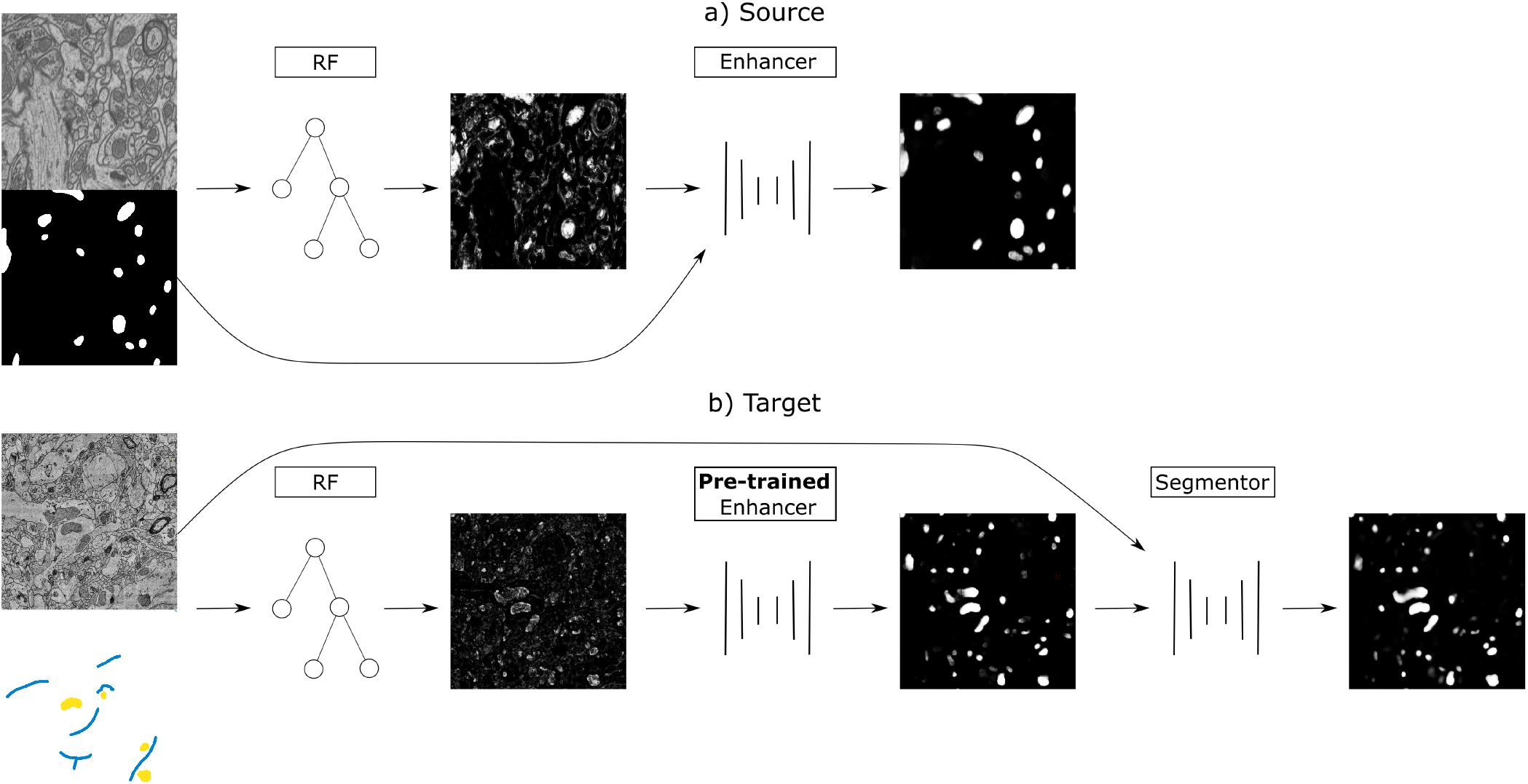
a) Training on the source dataset: many Random Forests are trained by subsampling patches of raw data and dense groundtruth segmentation. Random Forest predictions are used as inputs and groundtruth segmentation as labels to train the Prediction Enhancer CNN to improve RF segmentations. b) Domain adaptation to the target dataset: a RF is trained interactively with brushstroke labels. The pre-trained PE is applied to improve the RF predictions. Optionally, PE predictions are used as pseudo-labels to train a segmentation network for even better results with no additional annotations, but using a larger computational budget.

## 2 METHODS

Our approach combines the advantages of feature-based and end-to-end segmentation methods by training a Prediction Enhancer network to predict one from the other. On the target dataset, retraining can be limited to the feature-based classifier as its predictions – unlike the raw data – do not exhibit a significant domain shift if the same semantic classes are being segmented. In more detail, we propose the following sequence of steps (see also Figure 1):

1. Create training data for the Prediction Enhancer CNN by training multiple Random Forests on random samples of the densely labeled source domain.
2. Train the Prediction Enhancer using the RF predictions as input and the ground-truth segmentation as labels.
3. Train a Random Forest on the target dataset with a few brushstroke labels and use the pre-trained Prediction Enhancer to improve the predictions.
4. Use the improved predictions as pseudo-labels to train a CNN on the target dataset. This step is optional and trades improved quality for the computational cost of training a CNN from scratch.

Note that the Prediction Enhancer only takes the predictions of the Random Forest as input. Neither raw data nor labels of the source dataset are needed to apply it to new data. Our method can therefore be classified as *source-free domain adaption*, but the additional feature-based learning step allows us to avoid training set estimation or reconstruction, commonly used in other source-free or knowledge distillation-based approaches like Liu et al. (2021); Du et al. (2021). At the same time, we can fully profit from all advances in the field of pseudo-label rectification (Zhang et al. (2021); Prabhu et al. (2021); Wu et al. (2021); Zhao et al. (2021)), applying those to pseudo-labels generated by the PE network.

### 2.1 Prediction Enhancer

The Prediction Enhancer is based on the U-Net architecture (Ronneberger et al. (2015)). To create training data, we train multiple Random Forests on the dense labels of the source domain, using the same pixel features as in the ilastik pixel classification workflow (Berg et al. (2019)). To obtain a diverse set of shallow classifiers we sample patches of various size and train a classifier for each patch based on the raw data and dense labels. Typically, we train 500 to 1000 different classifiers. Next, we train the U-Net following the standard approach for semantic segmentation, using Random Forest predictions (but not the raw data) as input and the provided dense labels of the source domain as the groundtruth. To create more variability, we sample from all previously trained classifiers. We use either the binary cross entropy or the Dice score as loss function.

Segmentation of a new dataset only requires training a single Random Forest; its predictions can directly be improved with the pre-trained Prediction Enhancer. Here, we use ilastik pixel classification workflow, which enables training a Random Forest interactively from brushstroke user annotations.

### 2.2 Further domain adaptation with pseudo-labels

The Prediction Enhancer can improve the segmentation results significantly, as shown in section 3. However, it relies only on the Random Forest predictions, and can thus not take intensity, texture or other raw image information into account. To make use of such information and further improve segmentation results, we can use the predictions of the Enhancer as pseudo-labels and train a segmentation U-Net on the target dataset. We use either Dice score or binary cross entropy as loss and make the following adjustments to the standard training procedure to enable training from noisy pseudo-labels:

- use the RF predictions as soft labels in range [0, 1] instead of hard labels in {0, 1}.
- Add a consistency loss term similar to (Tarvainen and Valpola (2017)) that compares the current predictions to the predictions of the network’s exponential moving average. See also subsubsection 2.2.1.
- Use a simple label rectification strategy to weight the per-pixel loss based on the prediction confidence. See also subsubsection 2.2.2.

The combined loss function is defined as

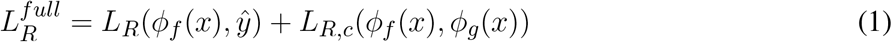

with consistency term *L*_*R,c*_ (see next section) and rectified pseudo labels *ŷ* (Equation 6). *L*_*R*_ denotes either binary cross entropy (*BCE*) or Dice loss (*dice*).

#### 2.2.1 Consistency Loss Term

For training with pseudo-labels we introduce a consistency term in the loss function, which is based on the “Mean Teacher” training procedure for semi-supervised classification Tarvainen and Valpola (2017). This method adds a loss term between the prediction of the network and its exponential moving average (EMA) to promote more consistent predictions across training iterations. We make use of this method for training a segmentation network *ϕ*_*f*_ with parameters *θ*_*f*_ from pseudo-labels. Its EMA is *ϕ*_*g*_ parametrized by

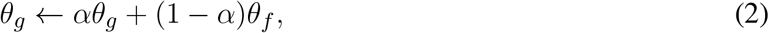

where we set the smoothing coefficient *α* to 0.999 following Tarvainen and Valpola (2017).

Given that we are comparing the per pixel predictions of the current network and its EMA, we use the loss function that is also employed for comparing to the pseudo labels: we either use the Dice loss

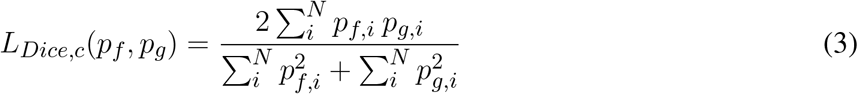

or the binary cross entropy loss

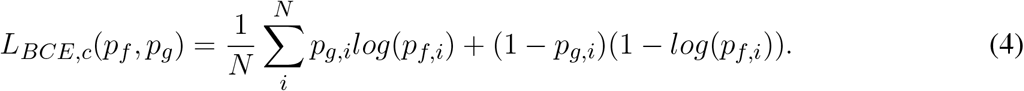

Where *x* denotes the input image, *p*_*f*_ = *ϕ*_*f*_ (*x*), *p*_*g*_ = *ϕ*_*g*_(*x*) and *N* is the number of pixels. The combined loss function is

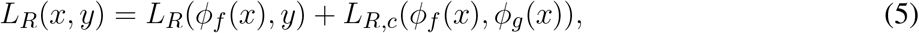

with pseudo-labels *y* and *R* either *Dice* or *BCE*.

#### 2.2.2 Label Rectification

Label rectification is a common strategy in self-learning based domain adaptation methods, where predictions from the source model are used as pseudo-labels on the target domain. Rectification is then used to correct for the label noise. Several strategies have been proposed, for example based on the distance to class prototypes in the feature space (Zhang et al. (2021)) or prediction confidence after several rounds of dropout (Wu et al. (2021)).

Here, we adopt a simple label rectification strategy based on the prediction confidence to weight the pseudo-labels *y* (which correspond to the predictions of the enhancer):

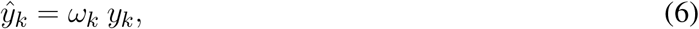

where *k* is the class index. For the case of foreground/background segmentation *k* ∈ {0, 1} and we define the per-pixel weight for the foreground class as

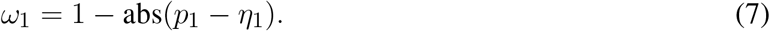

Here, *p*_1_ is the foreground probability predicted by the segmentation network and *η*_1_ the exponentially weighted average of foreground predictions:

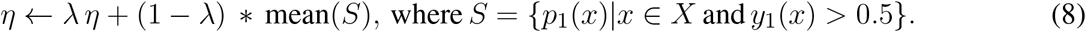

Here, *X* is the set of all pixels in the current batch and we set *λ* = 0.999 in all experiments. The weight *ω*_0_ for the background class is computed in the same manner.

## 3 RESULTS

### 3.1 Data & Setup

We evaluate the proposed domain adaptation method on challenging semantic segmentation problems, including mitochondria segmentation in EM, membrane segmentation in EM and LM as well as nucleus segmentation in LM. Table 1 summarizes all datasets used for the experiments.

**Table 1.** datasets used in the experiments

| Name | EPFL | VNC | MitoEM-R | MitoEM-H | Kasthuri | CREMI |
| --- | --- | --- | --- | --- | --- | --- |
| Organism/Tissue | Mouse/Hippocampus | Fruitfly/ventral nerve cord | Rat/cortex | Human/cortex | Mouse/cortex | Fruitfly/Brain |
| Modality | FIBSEM | ssTEM | sbEM | sbEM | ssTEM | ssTEM |
| Tasks | Mitochondria | Mitochondria, Membranes | Mitochondria | Mitochondria | Mitochondria | Membranes |
| Resolution | 5×5×5 nm | 45×5×5 nm | 30×8×8 nm | 30×8×8 nm | 30×3×3 nm | 40×4×4 nm |
| Reference | Lucchi et al. (2013) | Gerhard et al. (2013) | Wei et al. (2020) | Wei et al. (2020) | Kasthuri et al. (2015) | cremi.org |

(a) Electron Microscopy datasets used in the experiments.
| Name | Root | Ovules | DSB-FL | Monuseg |
| --- | --- | --- | --- | --- |
| Organism/Tissue | Arabidopsis/Lateral root | Arabidopsis/ovules | Various/nuclear stain | Human/kidney |
| Modality | Lightsheet | Confocal | Fluorescence | Histopathology |
| Tasks | Membranes | Membranes | Nuclei | Nuclei |
| Resolution | 0.25×0.1625×0.1625 μm | 0.235×0.075×0.075 μm | | |
| Reference | Wolny et al. (2020) | Wolny et al. (2020) | Caicedo et al. (2019) | Kumar et al. (2019) |

(b) Light Microscopy datasets used in the experiments.

Some of the datasets we use represent image stacks and could be processed as 3D volumes with different levels of anisotropy. We choose to process them as independent 2D images instead to enable a wider set of source/target domain pairs. If not noted otherwise, training from pseudo-labels is performed using the consistency loss term and label rectification (Equation 1). We use a 2D U-Net architecture (Ronneberger et al. (2015)) with 64 features in the initial layer, 4 downsampling/upsampling levels and double the number of features per level for all networks. The network and training code is based on the PyTorch implementation from Wolny et al. (2020). For all training runs we use the Adam optimizer with initial learning rate of 0.0002, weight decay of 0.00001. Furthermore, we decrease the learning rate by a factor of 0.2 if the validation metric is not improving for a dataset dependent number of iterations. We use binary cross entropy as a loss function for the mitochondria (subsection 3.2) and nucleus (subsection 3.4) segmentation and dice loss for the membrane segmentation (subsection 3.3).

### 3.2 Mitochondria segmentation

We first perform mitochondria segmentation in EM. We train the Prediction Enhancer on the EPFL dataset (the only FIB/SEM dataset in the collection) and then perform source-free domain adaptation on the VNC, MitoEM-R, MitoEM-H and Kasthuri datasets. For domain adaption, the Random Forest for initial target prediction is trained interactively in ilastik using a separate train split. The RF predictions are then improved by the PE and the improved predictions are used to as pseudo-labels for a U-Net trained from scratch (Pseudo-label Net). We compare to direct predictions of a U-Net trained for Mitochondria segmentation on the source domain EPFL (Source Net) and to the Y-Net (Roels et al. (2019)), a different method for domain adaptation, which is unsupervised on the target domain, but not source-free. We also indicate the performance of a U-Net trained on the target dataset as an estimate of the upper bound of the achievable performance (a separate train split is used).

Table 2 summarizes the resulting F1 scores (higher is better) for the source dataset and all target datasets. The Enhancer improves the Random Forest predictions significantly on all target datasets and the CNN trained from pseudo-labels further improves the results. The pseudo-label CNN always performs better than the source network or the Y-Net, which fails completely for the Kasthuri dataset where the domain gap is particularly large. Figure 2 shows an example of the improvements from RF to PE and PE to Pseudo-label Net.

**Table 2.** Results for mitochondria segmentation in EM. Quality is measured by the F1-score of the mitochondria prediction (higher is better). EPFL dataset is used as the source for domain adaptation by the Y-Net, Prediction Enhancer (PE) and Pseudo-label Net.

| Model / Dataset | EPFL | VNC | MitoEM-R | MitoEM-H | Kasthuri |
| --- | --- | --- | --- | --- | --- |
| Source Net | 0.933 | 0.695 | 0.738 | 0.591 | 0.723 |
| Y-Net | - | 0.713 | 0.781 | 0.678 | 0.0 |
| RF | 0.625 | 0.647 | 0.511 | 0.338 | 0.590 |
| PE | 0.824 | 0.840 | 0.705 | 0.624 | 0.778 |
| Pseudo-label Net | - | **0.884** | **0.793** | **0.751** | **0.834** |
| Target Net | 0.933 | 0.891 | 0.939 | 0.920 | 0.942 |

**Figure 2.**
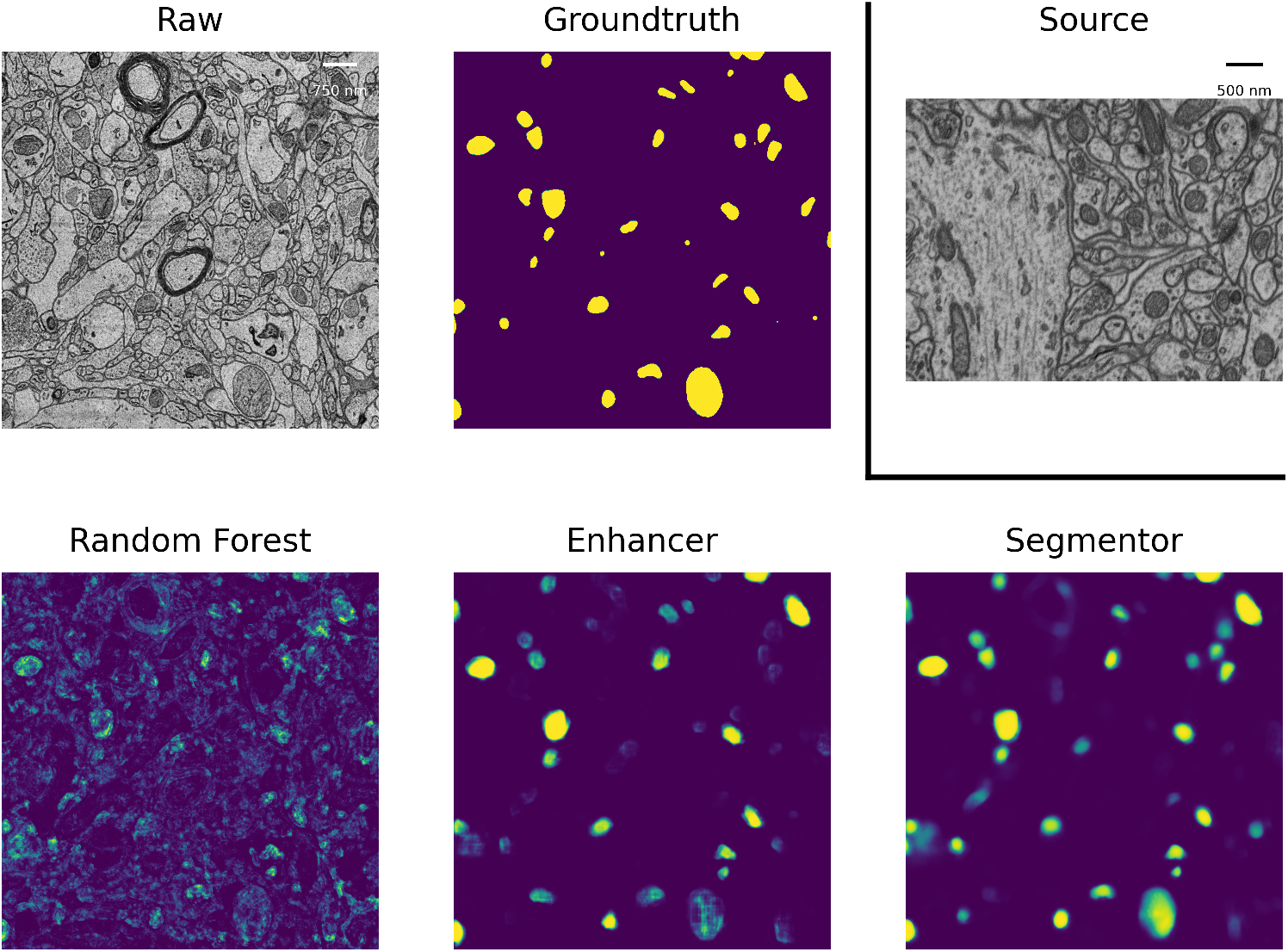
Mitochondria predictions of the Random Forest trained in ilastik, Prediction Enhancer and Pseudo-label CNN (“Segmentor”) as well as the groundtruth segmentation, on the MitoEM-H dataset. The Enhancer was pre-trained on the EPFL dataset; EPFL raw data shown under Source.

For the mitochondria segmentation task we also check if training the PE on multiple source datasets improves results. Table 3 shows that this is indeed the case, especially for the Kasthuri dataset.

**Table 3.**
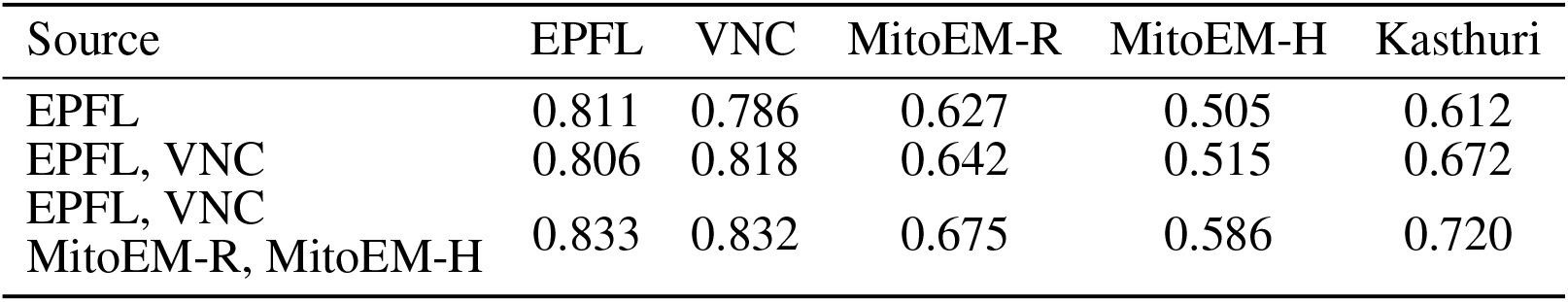
Mitochondria segmentation results for PE trained on multiple source datasets. The left column indicates the source datasets, quality is measured with the F1 score.

| Source | EPFL | VNC | MitoEM-R | MitoEM-H | Kasthuri |
| --- | --- | --- | --- | --- | --- |
| EPFL | 0.811 | 0.786 | 0.627 | 0.505 | 0.612 |
| EPFL, VNC | 0.806 | 0.818 | 0.642 | 0.515 | 0.672 |
| EPFL, VNC<br>MitoEM-R, MitoEM-H | 0.833 | 0.832 | 0.675 | 0.586 | 0.720 |

### 3.3 Membrane segmentation

We perform membrane segmentation both in EM and LM data. To evaluate membrane segmentations, we set up a Multicut based post-processing procedure following Beier et al. (2017) to transform the boundary segmentation into an instance segmentation, followed by evaluation with the Variation of Information (Meilă (2003)). We choose this more elaborate evaluation procedure as boundary segmentation is often used as the first step in instance segmentation pipelines and needs to be evaluated in this context. Simple evaluation by boundary F1 score is often not indicative of the actual quality of a boundary segmentation due to the large influence of seemingly small prediction errors, such as holes, on the follow-up instance segmentation. For the Variation of Information lower values correspond to a better segmentation.

In EM we perform boundary segmentation of brain tissue using the VNC dataset as source and three different datasets from the CREMI challenge (cremi.org) as target. Table 4 shows that the PE significantly improves the RF predictions for all three target datasets. The network trained on pseudo-labels can further improve results, especially for CREMI B and C, which pose a more challenging segmentation problem due to more irregular and elongated neurites compared to CREMI A. Both PE and Pseudo-label Net perform significantly better than a segmentation network trained on the source dataset. The segmentation results of a segmentation network trained on a separate split of the target dataset are shown to indicate an upper bound of the segmentation performance. Figure 3 shows the improvement brought by the PE and the Pseudo-label Net on an image from CREMI C.

**Table 4.** Results for boundary segmentation in EM. Quality is measured by the Variation of Information (lower is better) after instance segmentation via Multicut post-processing. Source Net and PE are trained on the VNC dataset and then applied to the three target datasets CREMI A, B and C. RF is trained interactively with ilastik on each target dataset.

| Model / Dataset | CREMI A | CREMI B | CREMI C |
| --- | --- | --- | --- |
| Source Net | 1.031 | 2.089 | 1.925 |
| RF | 1.092 | 2.231 | 2.363 |
| PE | 0.856 | 2.107 | 1.819 |
| Pseudo-Label Net | **0.840** | **1.806** | **1.582** |
| Target Net | 0.559 | 0.739 | 1.055 |

**Figure 3.**
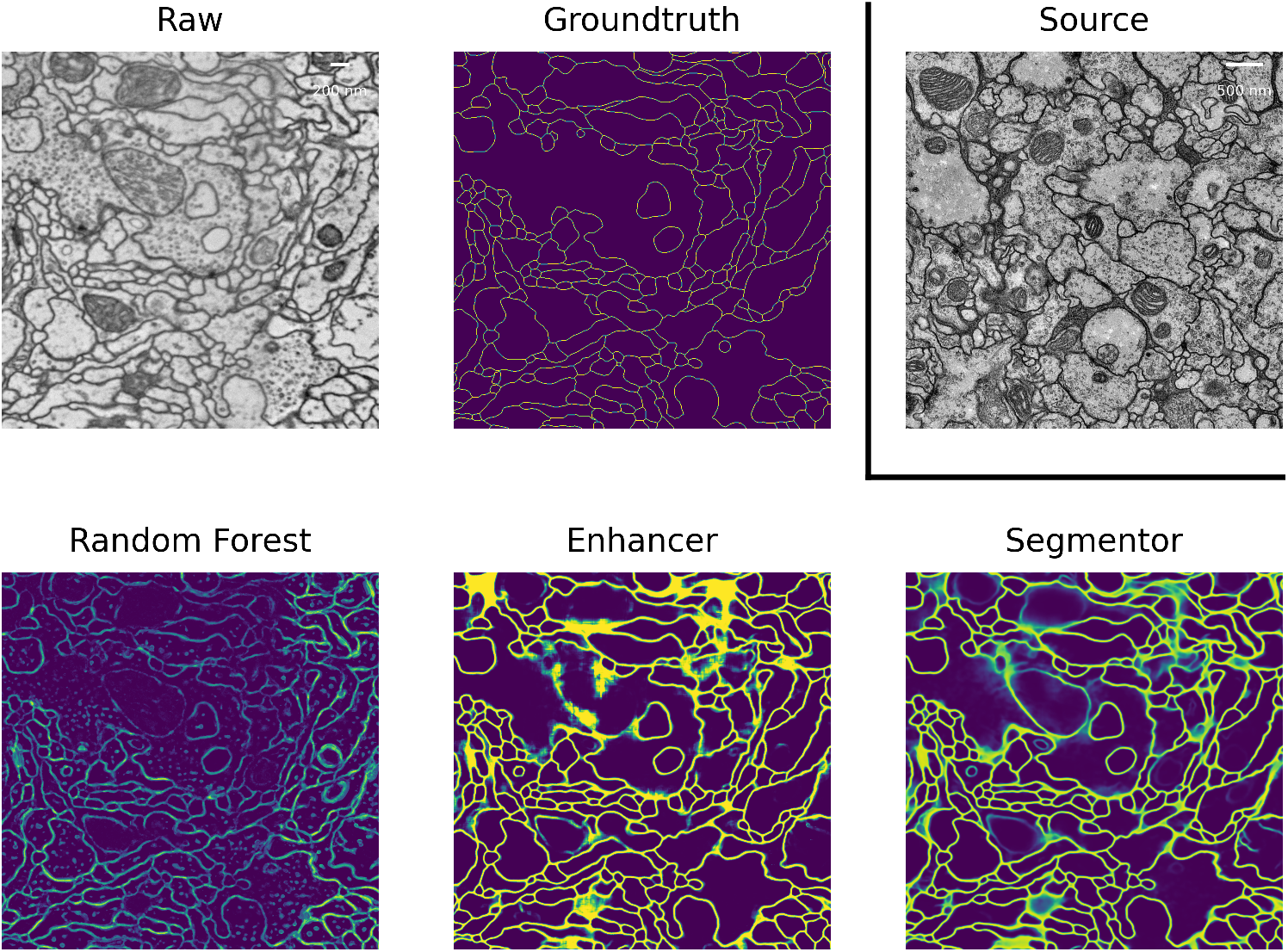
Boundary predictions of the Random Forest trained in ilastik, Prediction Enhancer and Pseudo-label Net, as well as the groundtruth segmentation, on the CREMI C dataset. The Enhancer was pre-trained on VNC, VNC raw data shown under Source.

In LM we perform boundary segmentation of cells in a confocal image stack of Arabidopsis ovules. We use a light-sheet image stack of Arabidopsis roots as source data. Note that we downsample both roots and ovules datasets by a factor of 2 for these experiments to increase the field of view of the segmentation networks. While this leads to source and target datasets with different resolutions (native resolution is 0.1625 *μ*m for roots and 0.075 *μ*m for ovules, see Table 1b) the size of the structure of interest matches best in this setting. Table 5 shows the results in the “Roots (LM)” column. While the PE improves the RF predictions and the Pseudo-label Net further improves result, we find that the source net performs better than any of our methods in this case. This can be explained by the fact that the quality of the RF segmentation is, in contrast to previous experiments, far inferior to the quality of the source network and the improvements afforded by PE and pseudo-label training are not sufficient to surpass the segmentation quality of the source network. Qualitatively, the RF predictions can be seen in Figure 4; the predictions amplify most of the signal in the image. This leads to a over-segmentation in the downstream instance segmentation, resulting in low quality segmentation. Note that the overall quality of results reported here is inferior compared to the results reported in Wolny et al. (2020). This can be explained by the fact that all models only receive 2D input, whereas the state-of-the-art uses 3D models.

**Table 5.** LM-Boundaries and cross modality experiments: Variation of Information after applying simple Multicut to the boundary predictions.

| Model / Source | Root (LM) | CREMI (EM) |
| --- | --- | --- |
| Source Net | **1.863** | 3.257 |
| RF | 2.442 | 2.442 |
| PE | 2.032 | 2.345 |
| Pseudo-Label Net | 1.982 | **2.309** |
| Target Net | 1.561 | 1.561 |

**Figure 4.**
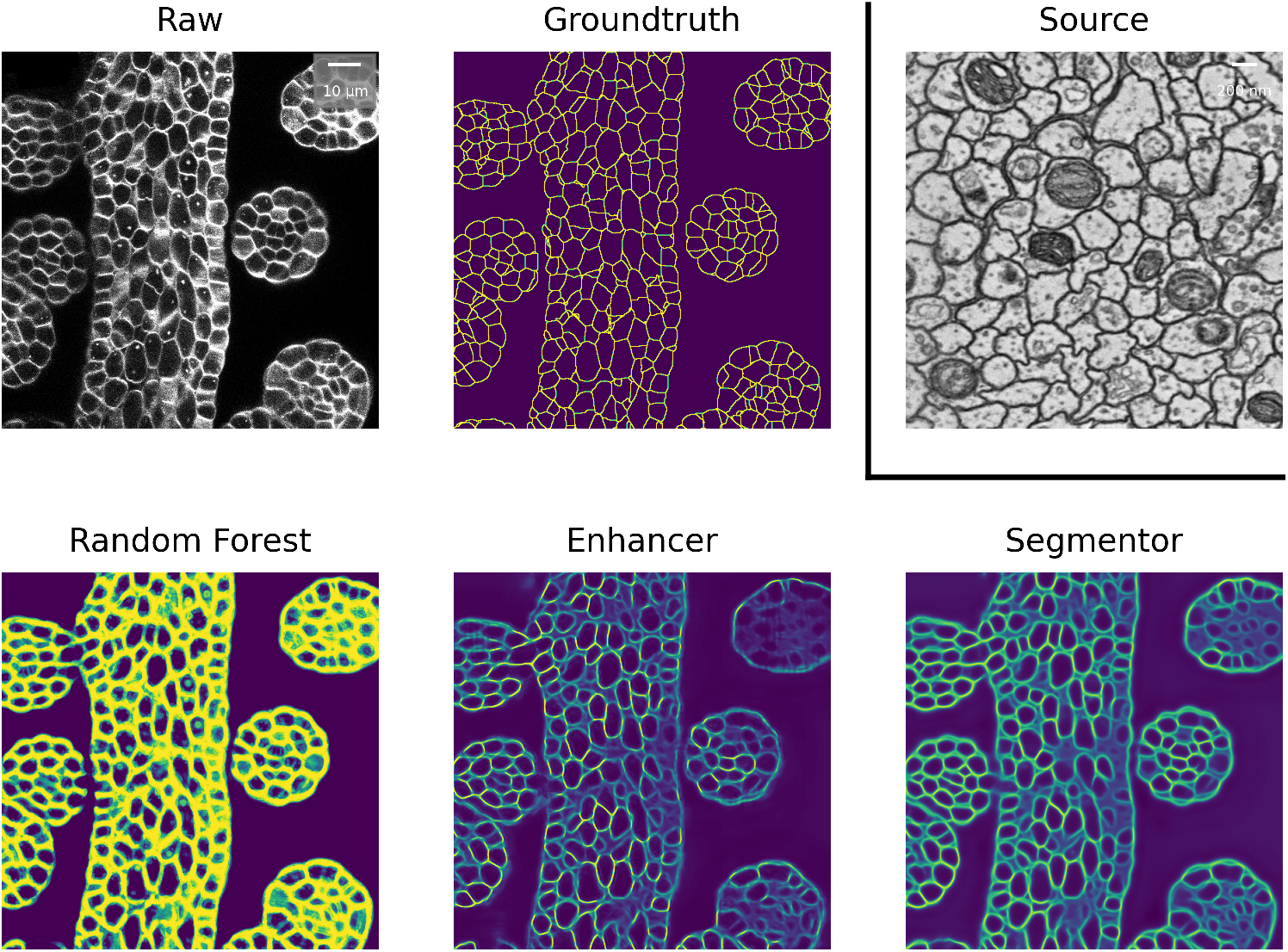
Boundary predictions of the Random Forest trained in ilastik, Prediction Enhancer and Pseudo-label Net, as well as groundtruth segmentation, on the ovules dataset. The enhancer was pre-trained on CREMI A, CREMI A raw data shown under Source.

We also experiment with a much larger domain shift and apply a PE that was trained on the EM dataset CREMI A as source. The results are shown in the “CREMI (EM)” column in Table 5. As expected, transfer of the source network fails, because it was trained on a completely different domain. However, the PE successfully improves RF predictions and pseudo-label training further improves the results. The fact that the PE only receives the RF predictions as input enables successful transfer in this case; while the image data distribution is very different in source and target domain, Random Forest probability maps look sufficiently similar. Furthermore, the resolution of the two domains differs by almost 3 orders of magnitude. However, the size of the structures in pixels is fairly similar, enabling successful domain adaptation. Figure 4 shows RF, PE and Pseudo-label Net predictions next to the source and target domain data.

### 3.4 Nuclei segmentation

As another example of cross-modality adaptation, we perform a an experiment for nucleus segmentation between fluorescence microscopy images from Caicedo et al. (2019) (DSB-FL) and histopathology images of the human kidney from Kumar et al. (2019) (Monuseg). Table 6 shows the results for using Monuseg as source and DSB-FL as target (column “DSB-FL”) and vice versa (column “Monuseg”). Here, the PE only affords a negligible improvement in the F1 score over the RF predictions but training from pseudo-labels improves the scores. We assume that the transfer of the PE is not very effective in this case because of very different nucleus sizes between the two datasets. The large domain shift is apparent from the fact that the Source Net does not generalize to the target domain at all in both cases.

**Table 6.**
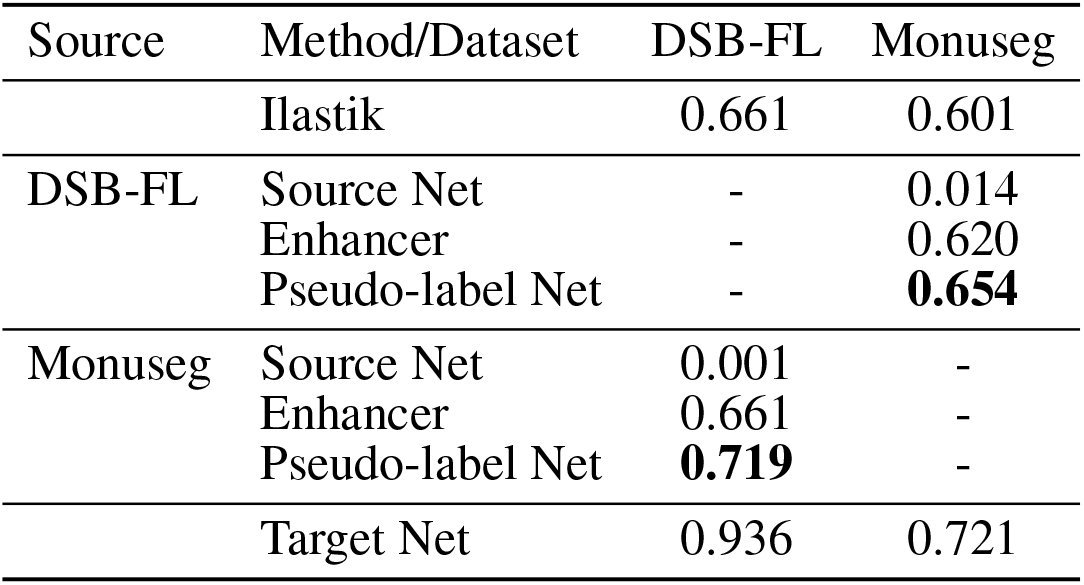
Results of nucleus segmentation. DSB-FL columns shows results for domain adaptation from Monuseg (Histopathology) to DSB-FL (Fluorescence), Monuseg column shows the opposite. The segmentation quality is measured by the F1 score.

| Source | Method/Dataset | DSB-FL | Monuseg |
| --- | --- | --- | --- |
|  | Ilastik | 0.661 | 0.601 |
| DSB-FL | Source Net | - | 0.014 |
|  | Enhancer | - | 0.620 |
|  | Pseudo-label Net | - | **0.654** |
| Monuseg | Source Net | 0.001 | - |
|  | Enhancer | 0.661 | - |
|  | Pseudo-label Net | **0.719** | - |
|  | Target Net | 0.936 | 0.721 |

### 3.5 Ablation studies

In the following, we perform ablation studies to determine the impact of some of our design choices on the overall performance of the method.

First, we investigate if the consistency loss (CL, equation 4) and label rectification (LR, equation 6) improve the accuracy obtained after pseudo-label training. We perform pseudo-label training for mitochondria segmentation on the VNC and MitoEM-R datasets using the PE trained on VNC to generate the pseudo-labels. We perform the training without any modification of the loss, adding only CL, adding only LR and adding both CL and LR. The results in Table 7 show that both CL and LR improve performance on their own. Combining them leads to an additional small improvement on VNC and to a slight decrease in quality on MitoEM-R.

**Table 7.**
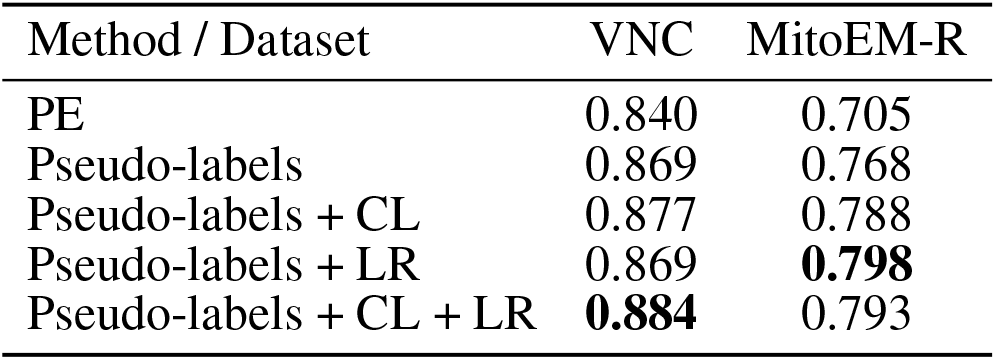
Results of pseudo-label network training using different loss functions. Mitochondria segmentation with EPFL as source dataset and VNC, MitoEM-R as target datasets. Segmentation accuracy is measured by the F1 score.

| Method / Dataset | VNC | MitoEM-R |
| --- | --- | --- |
| PE | 0.840 | 0.705 |
| Pseudo-labels | 0.869 | 0.768 |
| Pseudo-labels + CL | 0.877 | 0.788 |
| Pseudo-labels + LR | 0.869 | **0.798** |
| Pseudo-labels + CL + LR | **0.884** | 0.793 |

Using the same experiment setup, we also investigate whether using the PE enhancer for generating the pseudo-labels is actually beneficial compared to using the RF trained on target or using the source network. Table 8 shows that using the PE for pseudo-label generation significantly improves over the two other approaches.

**Table 8.** Results of pseudo-label network training using RF, Source Network and PE for label generation. Mitochondria segmentation with EPFL as source dataset and VNC, MitoEM-R as target datasets. Segmentation quality is measured by the F1 score.

| Pseudo-labels | VNC | MitoEM-R |
| --- | --- | --- |
| RF | 0.546 | 0.648 |
| RF w/ CL + LR | 0.584 | 0.656 |
| Source Net | 0.707 | 0.754 |
| Source Net w/ CL + LR | 0.794 | 0.765 |
| PE | 0.869 | 0.768 |
| PE w/ CL + LR | **0.884** | **0.793** |

### 3.6 Limitations

The high number of layers, their interconnections and especially skip-connections between them allow the U-net to implicitly learn a strong shape prior for the objects of interest. This effect is exacerbated in our Prediction Enhancer network as it by design does not observe the raw pixel properties and has to exploit shape cues even more than a regular segmentation U-net. While this effect is clearly advantageous for same-task transfer learning, it can lead to catastrophic network hallucinations if very differently shaped objects of interest need to be segmented in the target domain. To illustrate this point, we show the transfer of a PE learned for mitochondria on the EPFL dataset to predict boundaries on the VNC dataset and vice versa in Figure 5. The PE amplifies/hallucinates the structures it was trained on while suppressing all other signal in the prediction.

**Figure 5.**
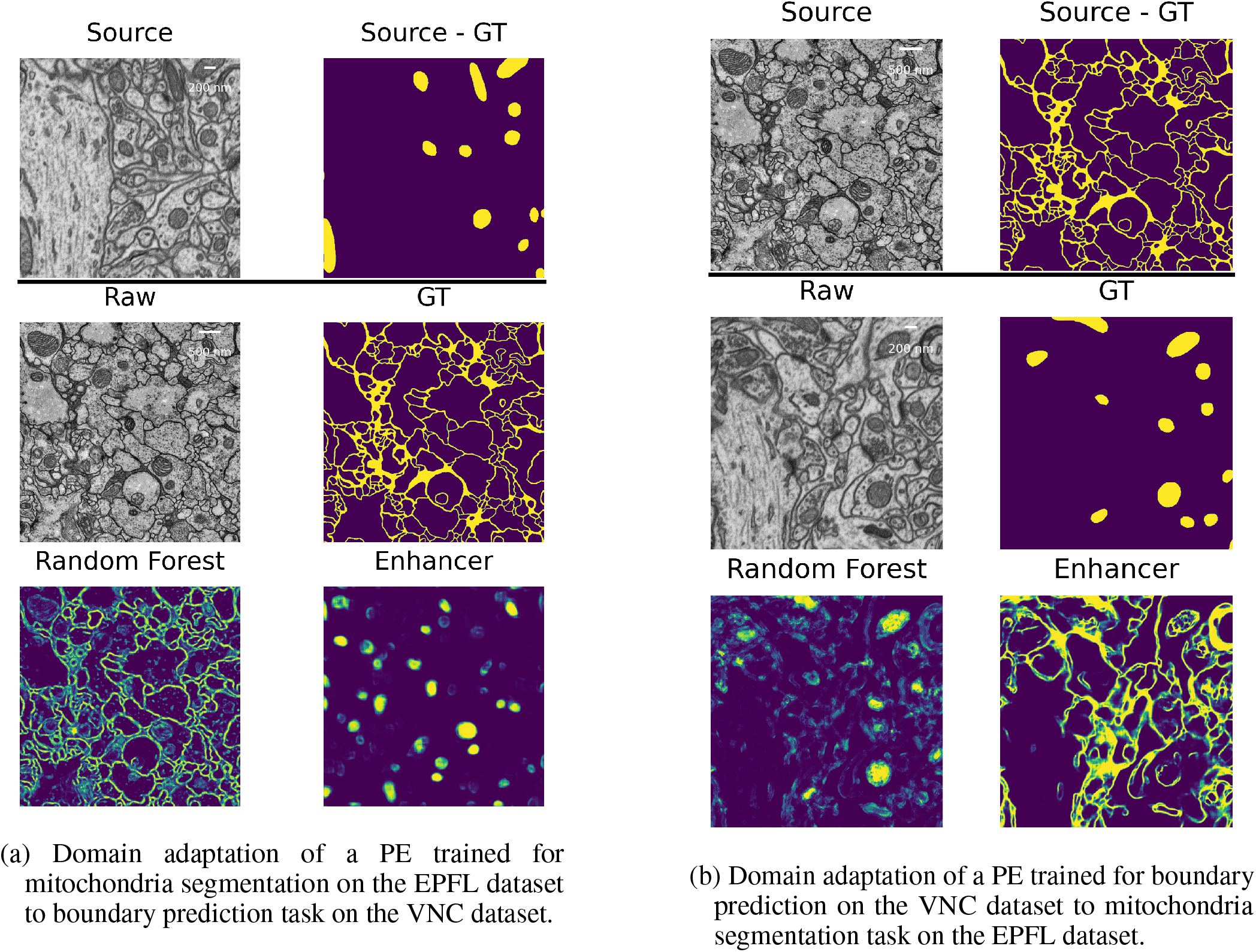
Failure case: different segmentation tasks in source and target datasets.

## 4 DISCUSSION

We have introduced a simple, source-free, weakly supervised approach to transfer learning in microscopy which can overcome significant domain gaps and does not require adversarial training. In our setup, the feature-based classifier which is trained from sparse annotations on the target domain acts as an implicit domain adapter for the Prediction Enhancer network. The combination of the feature-based classifier and the prediction enhancer substantially outperforms the segmentation CNN trained on the source domain, with further improvement brought by an additional training step where the Enhancer predictions on the target dataset serve as pseudo-labels. Since the Enhancer network never sees the raw data as input, our method can perform transfer learning between domains of drastically different appearance, e.g. between light and electron microscopy images. By design, this kind of domain gap cannot be handled by unsupervised domain adaptation methods which rely on network feature or raw data alignment. Furthermore, even for small domain gaps and in presence of label rectification strategies, pseudo-labels produced by the Prediction Enhancer lead to much better segmentation CNNs than pseudo-labels of the source network. We expect these results to improve even further with the more advanced label rectification approaches which are now actively introduced in the field.

The major limitation of our approach is the dependency on the quality of the feature-based classifier predictions. We expect that in practice users will train it interactively on the target domain which already produces better results than “bulk” training: in our mitochondria segmentation experiments, also shown in Table 2, there was commonly a 1.5-2 fold improvement in F1-score between interactive ilastik training in the target domain and RF training in a script without seeing the data. In general, the performance of the Prediction Enhancer will lag behind the performance of a segmentation network trained directly on the raw data with dense groundtruth labels except for very easy problems that can be solved by the RF to 100% accuracy. In a way, the Random Forest acts as a lossy compression algorithm for the raw data, which reduces the discriminative power for the Enhancer. However, the pseudo-label training step can again compensate for the “compression” as it allows to train another network on the raw data of the target domain, with pseudo-labels for potentially very large amounts of unlabeled data.

For simplicity, and also to sample as many source/target pairs with full groundtruth as possible, we have only demonstrated results on 2D data, in a binary foreground/background classification setting. Extension to 3D is straightforward and would not require any changes in our method other than accounting for potentially different z resolution between source and target datasets. Extension to multi-class segmentation would only need a simple update to the pseudo-label training loss.

In future work, we envision integration of our approach with other pseudo-label training strategies. Furthermore, as pseudo-label training can largely be configured without target domain knowledge, we expect our method to be a prime candidate for user-facing tools which already include interactive feature-based classifier training.

## Author Contributions

AK, AM, AW and CP have conceptualized the method. AM has implemented the method and run the experiments under the supervision of AK, AW and CP. AM and CP have drafted the manuscript and AK, AW and CP have written the final manuscript.

## Acknowledgments

We thank the EMBL IT Services for their support.

## Data Availability Statement

All datasets used for experiments are publicly available and can be otbained from their orignal publications, see Table 1. The code and models produced in this paper will be made available upon acceptance.

